# Distributed and convergent representations of tools in human parietal and anterior temporal regions revealed by fMRI-RSA

**DOI:** 10.1101/2024.04.22.590618

**Authors:** Ryo Ishibashi, Gina Humphreys, Ajay Halai, Azumi Tanabe-Ishibashi, Nobuhiro Hagura, Matthew A. Lambon Ralph

## Abstract

Classical models of tool knowledge and use are centred on dorsal and ventral parietal pathways. Theories of semantic cognition implicate a “hub-and-spoke” network, centred on the anterior temporal lobe (ATL), that underpins all concepts including tools. Despite their prominence, the two theoretical frameworks have never been brought together and the large discrepancy in the functional neuroanatomy addressed. We undertook a multiple-regression Representational Similarity Analysis (RSA) of task fMRI data with four (motor action, broad function, mechanical function, object structure) feature-based models. The motor action model correlated with the activation patterns in bilateral superior parietal lobules (SPL), while the models of broad function and mechanical effect aligned with the activation patterns in bilateral ATLs. The object-structure model correlated with activation patterns in bilateral middle occipital gyri. The results also showed that the ventral ATL activation patterns corresponded simultaneously with all RDM models except object structure. Furthermore, a standard univariate analysis using tool-familiarity ratings for parametric modulation revealed that classical tool-network regions (frontal, inferior parietal, and posterior middle temporal cortices) were increasingly active as the tool familiarity reduced. These results demonstrate that parietal and ATL regions are both crucial and motivate a major extension and revision of the neuroanatomical framework for tool use.

**Significance Statement:** This study provides definitive evidence for convergent tool representation in human anterior temporal lobe (ATL), outside the traditionally focused parietal lobe as the critical centre for human tool-use ability. For many years the parietal lobe was considered crucial in recognizing and planning use of familiar objects, while regions in temporal lobe received little attention. Our advanced multi-voxel analysis with artefact-resistive fMRI scanning revealed that both non-motor (tool-function) and motor (kinematics for tool use) information convergently represented in the left ventral ATL, while showing other distributed regions encoding distinct types of tool information in anterior temporal and parietal regions. These findings highlight the ATL’s crucial role in tool representation and necessitate a significant expansion of neuroanatomical framework for human tool-use ability.

## INTRODUCTION

Using tools is a crucial human ability, supporting various daily activities. However, how diverse tool-related information is encoded in the brain remains debated. Two major frameworks emphasize different brain areas: traditional action recognition and production theories focus on the parietal lobe, while recent semantic processing models highlight anterior temporal regions.

Since the 19th century, studies of (apraxia: Jackson, 1866; Liepmann, 1900) have identified the parietal lobe as key for action recognition and production, a finding supported by contemporary neuropsychological, neuroimaging, and neurostimulation studies (Boronat et al., 2005; Canessa et al., 2008; Gross & Grossman, 2008; Ishibashi et al., 2011; Pelgrims et al., 2011). Following this historical focus on parietal lobe, contemporary theories suggest that the parietal lobe processes distinct types of tool information via two pathways. The *dorso-dorsal* pathway, running from the occipital to the superior parietal lobule, is considered to compute object-directed action kinematics, while the *ventro-dorsal* pathway, along the inferior parietal lobule, is believed to integrate tool function into actions (Binkofski & Buxbaum, 2013; Buxbaum & Kalénine, 2010; Buxbaum & Randerath, 2018).

The temporal lobe has received far less attention in tool-use research. However, growing evidence suggests that anterior temporal regions play a crucial role. Studies of semantic dementia (SD) over the last two decades have shown that as knowledge of tools deteriorates due to bilateral ATL atrophy, patients also lose the ability to demonstrate their use, suggesting a strong link between conceptual knowledge and action (Bozeat et al., 2003; Corbett et al., 2009, 2011). Similarly, stroke patients with ATL infarction exhibit deficits in tool-use pantomime, further indicating its role in motor planning (Goldenberg & Randerath, 2015). Additionally, transcranial electrical stimulation of the ATL leads to enhanced manipulation-based decisions for familiar tools (Ishibashi et al., 2018). Despite these findings, ATL’s role in tool processing has received little attention in neuroanatomical models of human tool use. This overlook of ATL parallels with the historical bias occurred in language research, where early frameworks based on stroke aphasia primarily emphasized Wernicke’s and Broca’s areas. Over the past two decades, studies of semantic dementia, neuroimaging, brain stimulation, and computational modelling have led to an expansion of language models to include anterior temporal regions (Ueno et al., 2014; Saur et al., 2008; Lambon Ralph et al., 2017; Mesulam, 2023; Shimotake et al., 2015). These studies gave rise to the *hub-and-spoke* framework for semantic cognition, in which the ventral ATL serves as a central hub integrating diverse sensory and motor information, allowing the formation of generalizable conceptual representations (Lambon Ralph et al., 2010; Jackson et al., 2021). Despite this shift in language research, tool-use models remain largely parietal-centric and have yet to incorporate ATL’s role in tool-related processes.

Driven by the significant progress of these studies, and for our motivation to bridge language and praxis research, we investigated neural representation of tools beyond the parietal cortex. Specifically, we examined whether anterior temporal regions contribute to encoding any type of tool information. Our study leveraged two methodological advances. First, multi-echo EPI techniques for fMRI improve sensitivity within the ATL, overcoming susceptibility artifacts that have historically limited imaging of this region (Halai et al., 2014; Poser et al., 2006). Second, multiple-regression Representational Similarity Analysis (RSA) allows the examination of local neural coding of different types of object-related information. Our findings reveal that tool praxis information is predominantly represented in the superior parietal lobule (SPL), while tool function is primarily encoded in the anterior temporal lobe. Furthermore, the ventral ATL, the *semantic hub* region (Lambon Ralph et al., 2017), showed integration of multiple tool-relevant features, including both function and praxis knowledge. These findings strongly challenge the dominant parietal-centric model of tool use by demonstrating ATL’s crucial role in tool representation. By extending the tool-use network beyond the parietal lobe, our results suggest that human tool processing should be reconsidered within a broader perspective, incorporating both motor and semantic contributions in temporo-parietal network.

## MATERIALS AND METHODS

### Materials

We based our selection of tool stimuli on the neuropsychological study by Bozeat et al. (2002). The study examined tool-relevant cognitive performance of patients with semantic dementia utilizing a set of 90 different tools. We collected visual photo images of the corresponding 90 tools and used them for the present experiment (see Supplementary Data). Each collected image was converted to a 24-bit bmp file and then scaled to fit into the same image dimensions (400 × 300 pixels) using Windows Paint software.

### Participants

We recruited 22 graduate/undergraduate students in the University of Manchester. Their age range was 18-30 years (Mean 21.9; SD = 3.2). All had normal or corrected-to-normal vision and were right-handed (mean score in Edinburgh Handedness Inventory was +89.0). They gave their informed consent before taking part in this study and were reimbursed for their time. Two of them had to cancel their participation during the experiment (one at their will, another for a technical problem with the MRI scanner). Data from another participant had irreversible error (mixed-up slices) detected right after the recording. We therefore used data from the other 19 participants in the following analyses.

### Experimental Design

#### Procedure

Participants were engaged in visual recognition of tools while lying supine in the MRI scanner (Fig.1). The stimuli were presented with a projector system and seen in a mirror attached to the head coil. To make sure that the participants attend to the meaning of each stimulus, they were asked to judge whether the tool is used indoors or outdoors, and reported their decisions by pressing a button on a response box in their right hand. The responses were made using their right index and middle fingers. The assignment of buttons to the two (indoor/outdoor) categories was counterbalanced across participants. Each of the 90 stimuli appeared only once in a single continuous scanning (run) and stayed on the screen for 1.2 seconds regardless of their responses. A fixation cross on a grey background was presented when no stimulus is presented. The interval time between stimulus presentations were determined in advance using Optseq program (Dale, 1999; https://surfer.nmr.mgh.harvard.edu/optseq/) for each run. The intervals were set to range between 3.5-9.3 seconds by the minimum steps of 250ms and to have the mean of 4.8 seconds. The average stimulus onset asymmetry (SoA) was hence 6.0 seconds. A 16-second interval was added at the beginning and the end of each run to accommodate extra margins for the baseline. We used E-Prime software (Psychology Software Tools inc.) to implement the presentation of stimuli and the recording of the participants’ responses.

**Figure 1.**
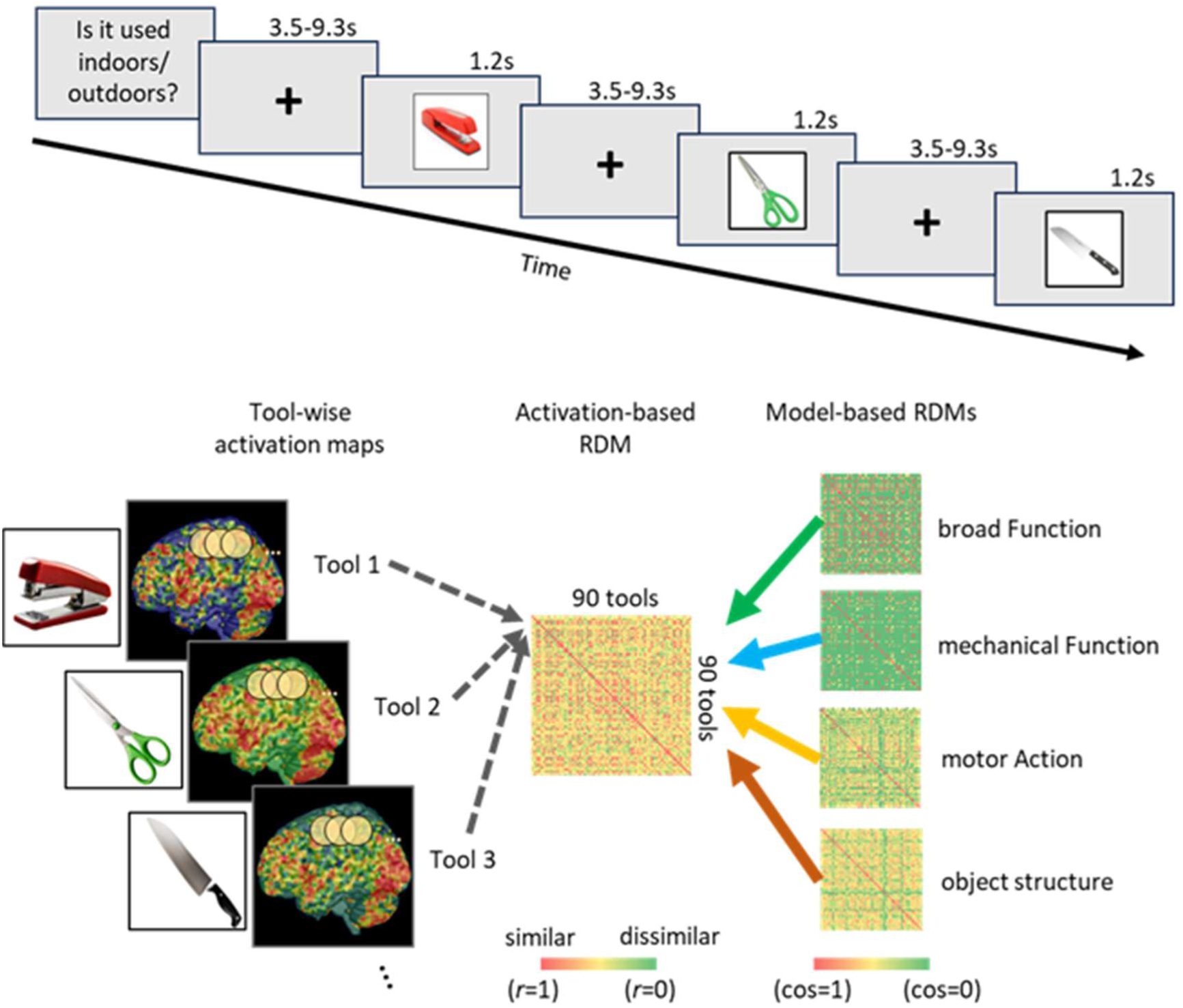
Top) Schematic of the behavioural task within the scanner; Bottom) Overview of a multiple-regression RSA for a given searchlight. The estimated standardized regression coefficients were registered for the location (centre) of the searchlight. Note that the actual analysis was conducted on the combined activation-based RDMs from all the participants. This mixed-effect regression analysis was repeated moving the searchlight across the whole brain to yield the coefficient map for each model.

Participants performed the task in 5 runs (approximately 55 minutes in total with in-scanner breaks between runs) with functional scanning, following an initial scanning of the structural T1 data of their brain (about 5 minutes). Before entering the scanner, they practised the task on a laptop PC. All stimuli appeared just once in the practice with the tool name shown below the image. This was to enhance their correct recognition of these images. After all runs in MRI scanner, the participants were asked to provide a familiarity rating for each tool on a Likert scale between 1 (“not at all”) and 5 (“very”). This rating task was conducted to confirm the familiarity of the tools across participants and to establish inclusion/exclusion of the tools in the main analysis: only tools with a familiarity rating>4.0 (on average) were included in the later RSA analysis in order to ensure that representational analyses were based on items that participants knew well. The rating resulted in 22 of 90 items having the mean familiarity <4.0. We used all items in a subsequent parametric-modulation analysis of the effect of familiarity in these task fMRI data (see below). The accuracy of the indoor/outdoor judgments on the included 68 items (58 indoor, 10 outdoor tools) was very high: 91.8% (SD=2.8). The mean response time was 749.9ms (SD=48.6).

### Image Acquisition with Dual-Echo fMRI sequence

We used a dual-echo sequence with 12ms and 35ms echo times for data acquisition (Halai et al., 2014; 2015). This method helps to reduce the effect of signal-dropouts at the basal parts of anterior temporal lobes and orbitofrontal cortex. Because tissues with heterogeneous magnetic susceptibility (e.g. cortex, bone, air) lie in close vicinity in these areas, canonical fMRI protocols with single echo time of around 35ms create distortions and signal loss in these regions. Given that signal dropout increases with echo time, the shorter of the dual echoes provides better signal recovery in these challenging areas but with less contrast, whilst the longer echo time retains contrast for other brain areas (Halai et al., 2014; 2015). Thus, we adopted this acquisition method for our experiment.

### Statistical Analysis

#### Preprocessing

The preprocessing steps followed the method established in previous dual-echo fMRI studies (Halai et al., 2014; 2015). The short-echo and long-echo images were re-aligned to the first volume in each run and then the signals were combined using a simple average at each voxel. The combined data were then passed to the subsequent pre-processing steps on the Statistical Parametric Mapping software (SPM8, https://www.fil.ion.ucl.ac.uk/spm/), which included slice-timing correction to the first (bottom) slice in each volume, coregistration of the first EPI volume with individual T1 image, and normalisation using DARTEL algorithm (Ashburner, 2007). No smoothing was applied to the data as the main purpose of this study is to explore multi-voxel patterns.

### Preparation for RSA: Univariate Analysis for Estimating Tool-Wise Activation Patterns

Voxel-wise GLM analysis was conducted using SPM8. We adopted a Condition-Rich design (Kriegeskorte et al., 2008) to model the BOLD signals at each voxel, setting each of the 90 stimulus as a different condition. Participants’ key-presses were modelled as an additional single ‘responses’ condition. Six volume-realignment parameters (6 parameters) from the preprocessing were also added to regress out head-motion. We also applied a high-pass filter of 128 seconds to remove low-frequency noise at each voxel. After estimating weights for all regressors, we calculated subject-wise contrast images for each of the 90 stimuli versus implicit rest. Excluding activation maps for the low-familiarity tools, a set of 68 × 19 contrast images were created as the target data for the following RSA analysis.

### Multi-Voxel RSA with Linear Mixed-Effect Regression

We adopted a multi-regression RSA approach (Kriegeskorte et al., 2008; Nili et al., 2014; Tucciarelli et al., 2019). Four models were used to represent different types of tool information: broad function (bF), mechanical function (mF), motor action (A), and object structure (Str). The models were created based on a normative object use feature study which was part of a previous study on investigation of tool knowledge in patients with semantic dementia (Bozeat et al., 2002). These models consisted of codes on the existence (1) or absence (0) of certain features on each of the 90 tool stimuli, providing a binary-code vector for each tool concept. The “broad function” of tools indicated the possible situations each tool can be used in. The “mechanical function” model (mF) involves a short description of mechanical purpose which each tool can be used for. The “motor action” model consists of the types of hand and arm movements involved in using each tool. The “object structure” model consists of visible structural properties of the tool. For each of the four models, cosine similarity was calculated for each pair of tools and then 1-cosine was used as the model’s representational dissimilarity matrix (RDM). The object structure model was involved in the analysis as a ‘control’ regressor. It was expected to prove the quality of the data and validity of the analytical procedure by showing high correlation in the occipital visual cortex.

We conducted both whole-brain and region-of-interest (ROI) analyses. In the whole brain analysis, we used a searchlight method with a linear mixed effect model (LME) analysis. The group-level mask image calculated from DARTEL processing was used as the area to explore. A spherical searchlight with 9 mm radius (involving 123 voxels) around each voxel was used to extract the multi-voxel patterns. Pearson’s correlation coefficients (*r*) across the tool stimuli were calculated and 1-*r* was used as the activity-based RDM. We conducted a mixed effect multiple-regression analysis with the upper triangular part of the activation-based RDM as the dependent variable and those of model RDMs as explanatory variables. Weighting of the four regressors were thus estimated for each searchlight location. This step was repeated for the search light across the whole brain. We created four T-value maps for the four model regressors, truncated them with uncorrected voxel-wise threshold of p<.001 and applied cluster-extent thresholding of p<.05 corrected for FWR to detect significant clusters. Only positive values were mapped and considered legitimate targets of significance tests in this RSA.

In the ROI analysis, we examined five anatomical ROIs in both hemispheres: two ROIs in parietal lobe (SPL, SMG), two in temporal lobe (ATL, pMTG), and one at the middle occipital gyrus (MOG). The parietal ROIs were chosen as the representative parts in the dorsal (dorso-dorsal and ventro-dorsal) pathways and the temporal ROIs as the representative parts along the ventral pathway. The MOG was chosen as a control region to examine the correspondence between low-level visual features (i.e., object-structure model) and the activation patterns in the visual cortex. We used WFU pick atlas (Maldjian et al., 2003) and AAL3 (Rolls et al., 2020) toolboxes to define four of the five ROIs. The ATL mask for a previous meta-analysis for temporal-lobe activation in fMRI studies (Rice et al., 2015) was utilised as the ATL-ROI. Regression analysis was conducted on the activation-based RDMs in each ROI. As in our searchlight analysis, only positive weights were considered targets of significance tests in this analysis.

Following these two analyses, we also conducted a conjunction analysis on the results of the models in the searchlight analysis. As we were interested in where the various semantic components are combined, the conjunction analysis was applied for the results of the first three models excluding the object structure (i.e., broad function, mechanical function and motor action) excluding the control model (object structure). The three T-maps were combined taking the minimum T statistics at each voxel. Significance tests were performed the same way as in the searchlight analysis. A voxel-wise threshold of α<.001 was first applied on each voxel (equivalent to p<.001 for the combined T map, i.e. p<.10 for each model). Cluster-extent threshold of p<.05 was then applied with FWE correction for the number of detected clusters.

### Univariate Analysis with Tool-Familiarity as Parametric Modulator

To examine the general task-related activation in this study and its correlation with the level of the participants’ tool knowledge (i.e., rated familiarity on the stimuli), we also conducted a univariate activation analysis on the same pre-processed data using each tool’s mean familiarity as a parametric modulator. All tool-presentation events were combined into a single “Stimulus” condition. Stimulus-wise mean familiarity values were added into the design matrix as a parametric modulator of the stimulus-evoked activation. The participants’ key-presses were also included in the model as a regressor of no interest. Both positive and negative contrasts were calculated for the base activation (task-positive/task-negative) maps and for the parametric-modulation (familiarity-/unfamiliarity-correlated) maps. The group-level activation maps were computed for each contrast and voxel-wise significance was tested using one-sample t test. Images were thresholded at a cluster-forming threshold of p<.001 and then small clusters were removed based on FWE-corrected cluster-extent threshold of p<.05. The results were compared with the results of our previous fMRI/meta-analysis studies reporting Semantic-/Executive-Control and Task-Negative networks (see also the caption of Fig.5).

## RESULTS

Fig.2 shows the results of the whole-brain searchlight analysis. The two function-knowledge models, broad-function (green) and mechanical-function (light blue), had high correlations mostly within the bilateral temporal lobes, with largest clusters at the anterior temporal lobes. The motor-action model (yellow) highlighted some parietal areas including bilateral superior parietal lobules and the right inferior parietal lobule. It also had high correlation with the activation patterns in the right inferior temporal gyrus and the left ATL. The object-structure model had high correlation in the bilateral occipital regions, roughly corresponding to the secondary visual cortex. The local maxima of the significant clusters for the four feature models are listed in Supplementary Tables 1.

**Figure 2.**
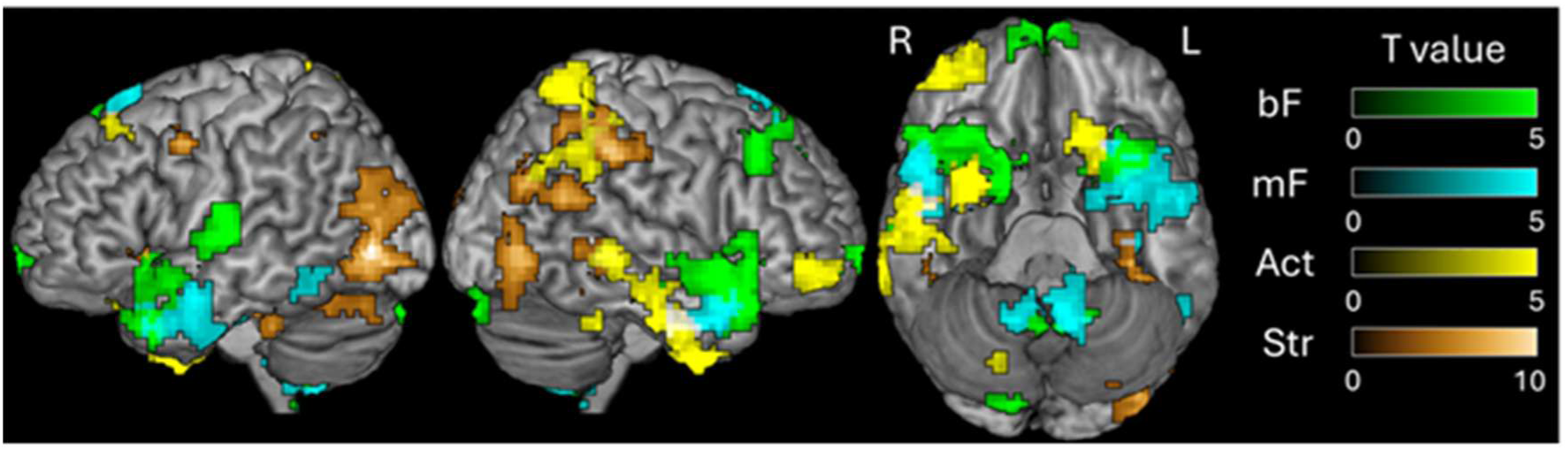
Results of the whole-brain searchlight analysis. Voxel-wise threshold was set at uncorrected p<0.001. Cluster-extent threshold was applied for each model map with an FWE-corrected threshold for p<.05. The minimum size of significant cluster was 77, 79, 84, and 78 voxels for the broad-function, mechanical-function, motor-action, and object-structure models, respectively. bF: broad function, mF: mechanical function, Act: motor action, Str: object structure.

The ROI analyses built on the searchlight results and allowed us to focus on key areas implicated in the classical praxis frameworks as well as the hub-and-spoke semantic model. Fig.3 show the results of the ROI analyses, dividing the ROIs by existence/absence of a significant effect of any of the four models. Fig.3-A shows that bilateral SPLs (involving the upper bank of IPS) had significant similarities with the motor-action model, the ATLs had correspondence with broad-/mechanical-function models, and the middle occipital gyri activates in high similarity with the object-structure model. The right SMG also had a significant correspondence with the structure model. The left SMG and the bilateral pMTGs, as indicated in Fig.3-B, had no significant similarity in their activation patterns with any of the four feature models.

**Figure 3.**
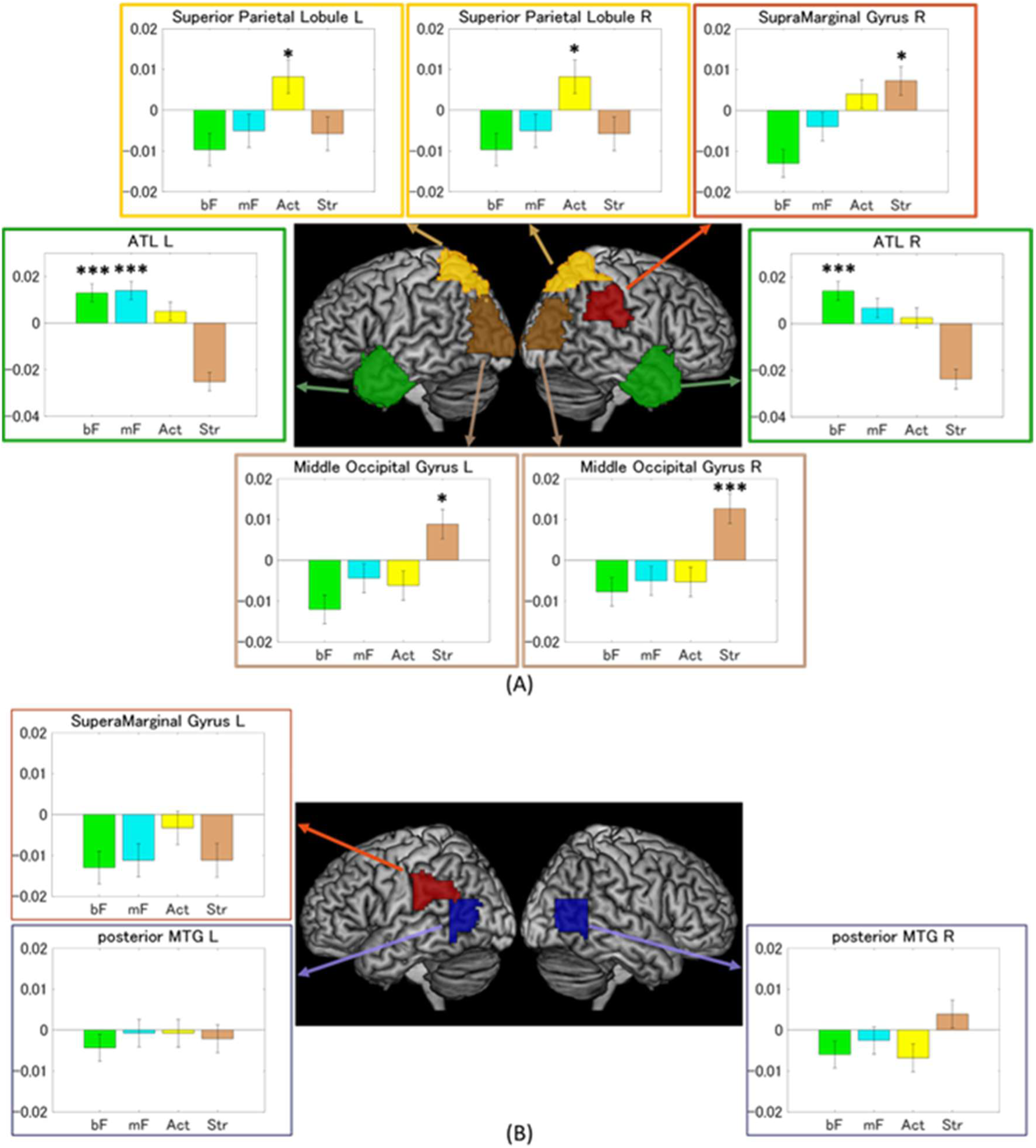
A) Regression coefficients for anatomical ROIs with significant positive effect by at least one of the four models. B) Regression coefficients for anatomical ROIs with no significant positive effect by any of the four models. (bF: broad function, mF: mechanical function, Act: motor action, Str: object structure. All ps>.05)

While the strongest contrast was observed between action and function models, the distribution of the significant clusters for each model gives important additional insights. Motor-action information was not exclusively represented in the parietal regions of the brain, but also represented in the bilateral temporal lobes. Some bilateral temporal areas also showed similarity to the motor-action model, making anterior parts of bilateral ATL the most probable overlapping areas across models. Accordingly, we conducted a conjunction analysis to confirm if any area represents both function and action types of tool knowledge (i.e., similarity to the first three models). Four clusters were found significant (Fig.4-A, Supplementary Table 2). The largest cluster locates within the left ATL extending into left inferior frontal gyrus (see Fig.4-B for the confirmation of regression results at the peak-ATL searchlight). The right ITG (covering the caudal ATL and fusiform gyrus), left insula and cerebellum were also detected as showing significantly similar activation patterns across the three tool-knowledge models.

**Figure 4.**
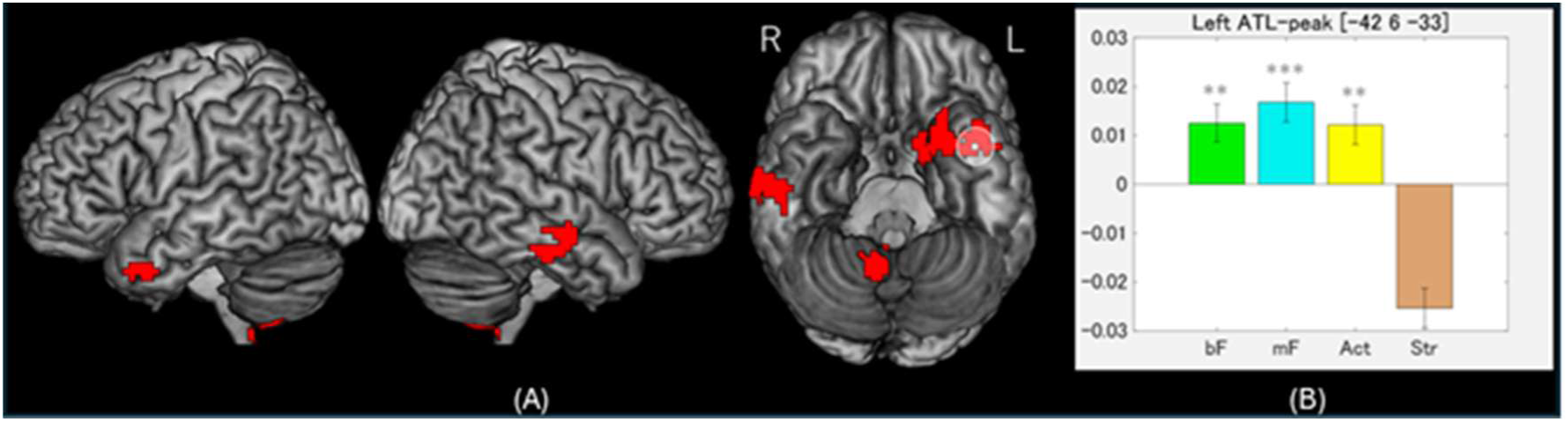
A) Areas with representational similarities to all three types of tool information. Cluster-extent threshold was applied for each cluster with the FWE-corrected threshold for p<.05. The minimum size of significant cluster was 60 voxels. The white dot and circle indicate the location of the voxel with the peak T value and its corresponding searchlight in the left ventral ATL. B). Estimated weights of the four regressors at the peak searchlight from the conjunction analysis. Grey stars at the top indicate the level of statistical significance. Note that significant tendency at the level of p<.10 is obvious for the first three regressors by the nature of this analysis. The bar plot and the statistical results are presented here for confirmatory purpose. ***: p<.001, **: p<.01. bF: broad function, mF: mechanical function, Act: motor action, Str: object structure.

### Functional-MRI Results: effects of tool familiarity across the tool-network activation

The univariate task-related activations (Fig. 5-A top row, left) revealed the typical fronto-parietal and occipito-temporal ‘tool-network’ areas found in previous studies and ALE meta-analyses (Chao & Martin. 2000; Federico et al., 2023; Ishibashi et al., 2016; Lewis 2006; Wang et al., 2018). These same areas were also observed in the map of areas where activation correlated with the degree of unfamiliarity with each tool (i.e., more activation for less familiar tools; Fig. 5A second row, left). As these same areas have been implicated in semantic-executive control, we compared them directly with the results from previous work on semantic and domain-general control (Humphreys et al., 2017). There was striking resemblance including the left pMTG/occipito-temporal cortex, SMG, and bilateral frontal/prefrontal regions (Fig. 5-B, left). The overlap in the posterior pMTG also corresponded to the location of common activation for semantic activation and tool cognition indicated by a recent large-scale meta-analysis (Hodgson et al., 2023; Fig.5-B in the enlarged view).

**Figure 5.**
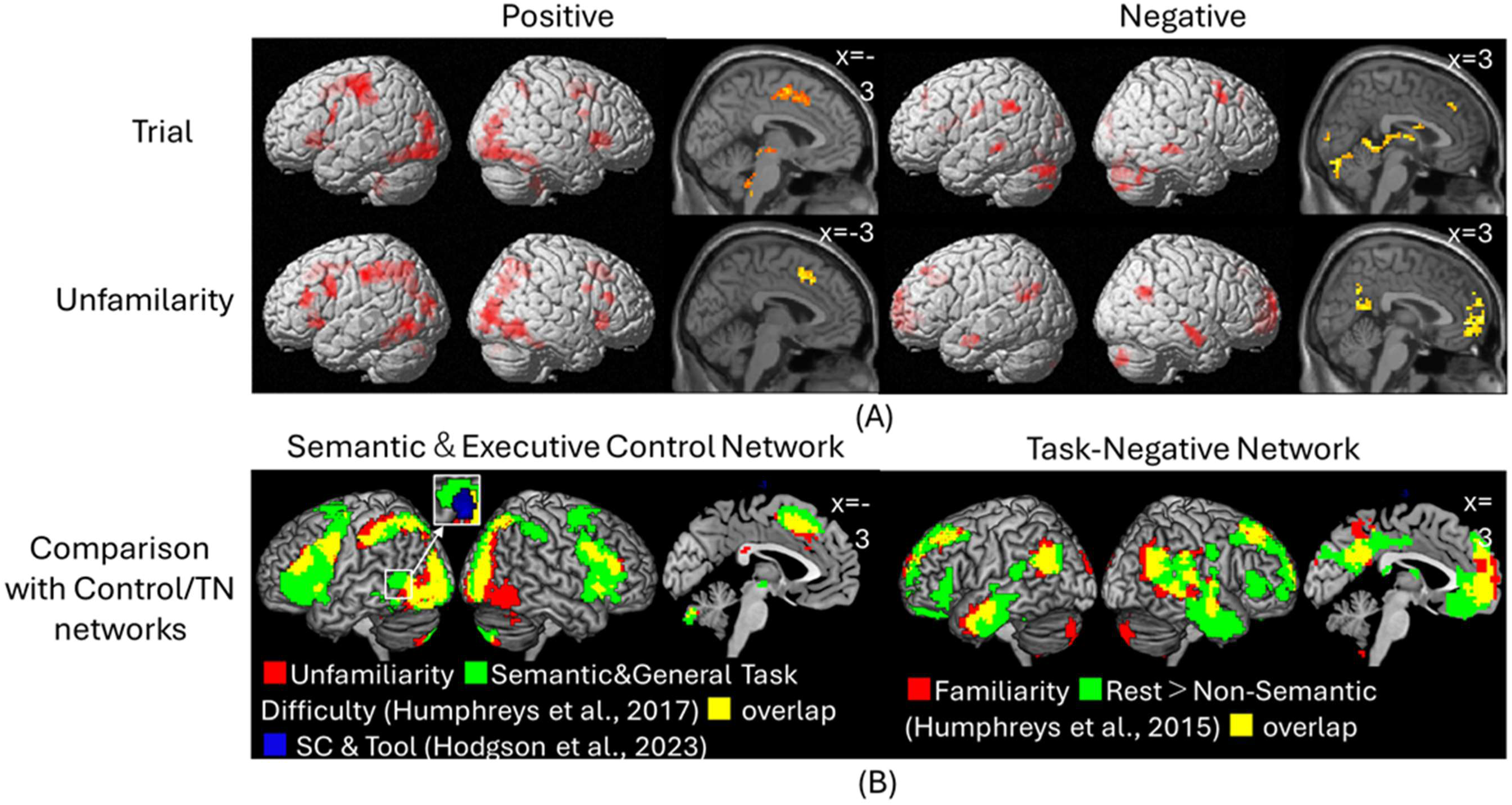
Mass-univariate activation maps. A) task-related base activation (top) and its parametric modulation by unfamiliarity of the presented tool (bottom); B) Overlays of the parametric modulation maps with other results from previous studies on general/semantic executive demands (left) and on the task-negative activation. Maps in panel A are all thresholded at a cluster-forming threshold of p<.001 and then small clusters were removed based on FWE-corrected cluster-extent threshold of p<.05. In panel B, the activation maps were thresholded at p<.05 for exploratory purpose. The Rest>Non-Semantic contrast was thresholded at p<.005 for its relatively large sample size (N=69). Clusters smaller than 25-voxel size are not shown for the sake of brevity. TN: Task-Negative.

The reverse contrasts indicated that the regions positively correlated with tool familiarity were the dorsal medial prefrontal cortex (mPFC), posterior cingulate cortex (PCC), and bilateral angular gyri (AG) ―typical task-negative areas which deactivate in various cognitive tasks, including the current fMRI task (Fig. 5-A right). There were also some familiarity-correlated activations in the lateral ATLs, which are also sometimes a subcomponent of the task-negative network (Buckner et al., 2008; 2019; Jackson et al., 2019; Menon et al., 2023) and tend to show enhanced deactivation in line with increasing task demands (Humphreys et al., 2015). In line with this result, there was a strong correspondence (Fig. 5-B, right) of the familiarity-positive activation correlation map from this study with the task-negative activation for non-semantic tasks observed in a previous multi-experiment study (Humphreys et al., 2015), including overlap in the medial prefrontal-parietal regions (mPFC, PCC), bilateral AG, and bilateral lateral ATLs.

## DISCUSSION

By utilising contemporary fMRI acquisition methods and analytics, the current study was able to investigate, for the first time, how multiple aspects of tool knowledge are simultaneously represented in the healthy brain. The results not only reinforce the importance of classical dorsal parietal regions in human praxis but also heavily implicate the ATL as a major representational hub for multiple aspects of tool knowledge, and suggest that a significant expansion of the classical model of object use is necessary. We consider below how the relationship of the current results to: (i) classical models of praxis in the dorsal parietal cortex; (ii) the striking multifaceted contributions of the ATL to tool knowledge and use; and (iii) the role of posterior lateral temporal and inferior parietal regions in object tool use.

### SPL and the dorso-dorsal pathway

Consistent with the classical dual-route theory, we found that SPL/IPS activation was correlated with the motor action RDM. This region is known as the classical where/how pathway in visual information processing, and is thought to perform planning of hand-object interactions from visual information (Goodale & Milner, 1992; Goodale et al. 2005). Cortical damage in SPL/IPS causes deficient object-directed actions including dysfunctional reaching and grasping (Binkofski et al., 1998; Cavina-Pratesi et al., 2013; Karnath & Perenin, 2005; Perenin & Vighetto, 1988). As the motor action RDM consists of features of the grip and arm movements for tool use, the present results suggest that body-movement for tool use is indeed calculated/represented in the bilateral SPL/IPS regions. Corroborating this view, a recent study found successful decoding of planned grasp type from activation in both right and left anterior IPS (Michalowski et al., 2022), and activity in bilateral IPS during planning tool-use has been found to increase with the complexity of manipulating each tool (Ras et al., 2022). Some clinical studies also suggest that the SPL contributes tool-manipulation knowledge, as SPL-damaged patients struggle to show canonical manipulation of common tools without agnosia on the tools (Valério et al., 2021; Peigneux et al., 2001).

### The importance of the ATL semantic hub

A striking aspect of the current study was the importance of the ATL in the representation not only of the two forms of tool function knowledge but also motor actions (in fact, only the visual form similarity did not engage the ATL). This finding of the ATL encoding of motor information is in a close alignment with a recent study by Knights and colleagues, which reports ‘hand-movement’ representation in ATL activity while the participants were making a grip on tools in MRI (Knights et al., 2022). Though the ATL is yet to be incorporated into human tool-use models, our results provide clear evidence for necessary integration; in the same way that the ATL has been integrated into contemporary models of language and semantics (Chen et al., 2017; Frisby et al., 2023; Lambon Ralph et al., 2017; Mesulam, 2023; Patterson et al., 2007). The absence of the ATLs from models of praxis (and previously of language) probably reflects the neuropsychological origins of the classical models, which were founded on the study of patients after MCA (middle-cerebral artery) stroke. The middle-to-inferior aspects of the ATLs are in the watershed between the MCA and PCA (posterior-cerebral artery)(Lambon Ralph et al., 2017; Phan et al., 2005; Zhao et al., 2020), and thus unlikely to be damaged after embolic stroke (and even less likely to produce bilateral ATL damage, which is the atrophy distribution in semantic dementia; Ding et al., 2020; Rouse et al., 2024). Also, past functional MRI studies with healthy participants often had to overlook ATL activation or exclude it from the field of view, due to the confounding susceptibility artefact (Poser et al., 2006; Visser et al., 2010). Applying the dual-echo method and appropriate analytical pipeline (Halai et al., 2014), this study achieved sufficient signal quality in the ATL, and succeeded to demonstrate the region’s clear encoding for multiple aspects of tool information.

The precise role of ATL in human tool-use behaviour is not evident from this study alone, but combining the present finding with other evidence suggests a critical contribution of the ventral ATL not just for the combination of tool function and action, but also for the stable retrieval of each component of tool information. The discovery of ATL involvement in tool-representation is consistent with the semantic hub-and-spoke model (Lambon Ralph, 2017). The framework posits that the bilateral ATLs serve as a modality-general ‘hub’, integrating multimodal information over time to build generalisable conceptual knowledge (Lambon Ralph et al., 2010; Jackson et al., 2021). The benefit of having such a ’hub’ region, in addition to other modality-specific sensory/motor areas, is demonstrated in computational modelling studies comparing a semantic system with and without a hub (Lambon Ralph et al., 2007; Rogers et al., 2004), which showed that a system with an intermediate ’hub’ layer performs much better in activating precise sets of sensory/motor features of a given concept than a system without a hub (sensory/motor layers only). Given the present finding of the multifaceted tool encoding in the left ventral ATL and its stereotactic correspondence with the ’semantic-hub’ region, it is arguable that the region not only represents functional and motor information simultaneously, but also provides a coherent pattern of activation to facilitate retrieval of function/motor information in other modality-specific regions. This view about the causal role of the ATL for tool use fits well with previous investigations using TMS that demonstrate the critical causality of ATL activation to the recognition of tools and their associated actions (Ishibashi et al., 2011; Pobric et al., 2010).

Neuroanatomically, ATL involvement in praxis may seem somewhat surprising given its distance from other tool-related cortical areas. Although the cortical distances between ATL and SPL are large (almost as large as they can be within the same hemisphere), they have direct white-matter connectivity potentially allowing fast, efficient communication and emergent functional interactions. Human and non-human primate tractography studies have found that the superior ATL and inferior parietal cortices are connected along the middle longitudinal fasciculus (Menjot de Champfleur et al., 2013), which extends up to the SPL (Wang et al., 2013; Markis et al., 2013). In parallel, the inferior longitudinal fasciculus (ILF) not only connects the temporal pole with occipital visual areas (Tusa & Ungerleider, 1985; Catani et al., 2003) but also branches at the ventral occipito-temporal area to ascend, medially, into the dorsal parietal cortex, in both non-human primates (Selzer & Panja 1984; Schmahmann & Pandya, 2009) and humans (Lang et al., 2017). Whether this ascending parietal branch should be regarded a part of ILF remain controversial (Mandonnet et al., 2018; Bullock et al., 2022), however it is quite possible that neurons in the SPL and ATL are connected directly via midLF, or through a single intermediating region at the occipito-temporal cortex.

### Temporoparietal and ventro-dorsal pathway contributions

The null RSA results for the left pMTG and SMG are potentially puzzling given that these regions are frequently noted in tool-use/recognition (univariate) fMRI studies, as well as being a key area of damage in ideational apraxia (Goldenberg & Spatt, 2009; Goldenberg & Randerath, 2015; Hodgson et al., 2023; Ishibashi et al., 2016). Our univariate analysis with the same data revealed two important findings: (1) both the pMTG and SMG were activated by the tool decision task, and (2) these two regions, along with the inferior frontal gyrus (IFG), responded more strongly to the least familiar tools. In keeping with these results, one possible explanation is that these regions (along with PFC) do not represent tool information per se but support semantic/executive control (Jackson et al., 2021; Jackson, 2021; Lambon Ralph et al., 2017); a conclusion supported by convergent evidence from fMRI, rTMS and neuropsychology (Hallam et al., 2016; Jefferies, 2013; Whiteney et al., 2011; 2012). The overlay of our activation results and previous meta-analyses/imaging experiments on the comprehensive cortical activity responding to the degree of semantic control clearly indicate the considerable alignment in the fronto-temporo-parietal (FTP) network (Fig. 5), which corresponds to other consistent reports of this tri-regional network in functional and structural brain connectivity analyses (Jung et al. 2017; Spreng et al., 2013). This striking similarity of the current univariate results and FTP-control network suggests that the ventro-dorsal and pMTG regions are in fact a part of cognitive control circuit required for task-appropriate retrieval of tool information. In fact, patients with damage to these brain regions not only show multimodal semantic control deficits (semantic aphasia: Jefferies & Lambon Ralph, 2006; Jefferies et al., 2008; Noonan et al., 2013; Thompson et al., 2022) but also present with ideational apraxia (Corbett et al., 2009a, 2009b; 2011; 2015), a condition in which patients are compromised in orderly retrieval of multiple movement components for an action (e.g., making tea using a teapot, kettle and teabag in appropriate steps).

The reason why tool cognition and semantic control align in these same regions (see Fig. 5; and also a recent large-scale ALE examination: Hodgson et al., 2023) is not fully understood. It is possible that some executive planning is required or automatically triggered in the recognition of tools. Multiple recent reviews on the neural mechanisms of human tool use argue that tool use is not simply the recall of movement information for each tool, but rather a compositional process for integrating relevant physical information of the tool and expected target objects for a desired goal state (technical-reasoning hypothesis; Jarry et al., 2013; Osiurak & Badets, 2016; Lesourd et al., 2021). Although this account has focussed on the role of left SMG, the required executive process for tool-use ‘reasoning’ is likely to require contributions from multiple multi-demand and semantic control areas beyond SMG alone.

### Conclusion

The adoption of RSA and multi-echo fMRI shows that the cortical areas implicated in human tool cognition and use go beyond the parietal-centric dorso-dorsal and ventro-dorsal pathways. A more comprehensive account is needed, in which the classical framework is extended to include the ATLs’ essential role in representation of multiple aspects of tool knowledge, plus the support of the FTP semantic-control network for controlled, goal-directed tool-use actions.

## Acknowledgments

This study was supported by JSPS Overseas Research Fellowship to R.I. (H27-612), an MRC Career Development Award to A.H. (MR/V031481), an MRC Programme grant to M.L.R (MR/R023883/1) and an intramural award (MC_UU_00005/18).

## Contributions

R.I., A.T., and M.L.R. conceived the project. G.H., A.H., and R.I. prepared experimental settings for data acquisition. R.I. conducted analyses and prepared the first draft for the paper with joint supervision by N.H. and M.L.R. All authors reviewed, edited, and commented on the draft.

## Notes on FWHM estimation

The FWHM for cluster-extent thresholding was estimated based on a simulation of spatial auto-correlation between neighbouring voxels. We utilised the ratio of shared number of voxels between two neighbouring searchlights in the current setting (94 of 123 voxels in a single searchlight). On a MATLAB-R2022b platform, we first generated random Gaussian noise in an array of 123 × 68 cells. For each array, 29 × 68 cells were updated by another Gaussian-noise simulation keeping 94 × 68 of them unchanged. The 68-by-68 RDM was calculated for the original and the updated arrays, and then the upper-triangle part was extracted as a vector from each. These steps were repeated 19 times to simulate the noise data from 19 virtual participants. Then, the vectors were concatenated to create a single error vector for the original and updated vectors, respectively. This was to simulate the number of datapoints for the current multiple-regression RSA. The correlation between the original and updated error vectors were then calculated as an index of simulated pure-noise-based autocorrelation between a pair of neighbouring searchlights. The entire above procedure was repeated 10,000 times to simulate distribution of correlation values in multiple pairs of searchlights, and the average *r*=0.761 (SD=0.0031) was taken as the theoretical spatial autocorrelation between neighbouring datapoints in the current whole-brain analysis. The in-house script for this simulation is available upon request to the corresponding author.

## Supplemental Dataset

This manuscript is accompanied by an excel file “SupplementaryData.xlsx”, containing the full tool-feature list and their binary coding for 90 items (from Bozeat et al., 2002).

